# Long-term molecular turnover of actin stress fibers revealed by advection-reaction analysis in fluorescence recovery after photobleaching

**DOI:** 10.1101/2021.06.19.449123

**Authors:** Takumi Saito, Daiki Matsunaga, Tsubasa S. Matsui, Shinji Deguchi

## Abstract

Fluorescence recovery after photobleaching (FRAP) is a versatile technique to evaluate the intracellular molecular exchange called turnover. Physicochemical models of FRAP typically consider the molecular diffusion and chemical reaction that simultaneously occur on a time scale of seconds to minutes. Particularly for long-term measurements, however, an advection effect can no longer be ignored, which transports the proteins in specific directions within the cells and accordingly shifts the spatial distribution of the local chemical equilibrium. Nevertheless, existing FRAP models have not considered the spatial shift, and as such, the turnover rate is often analyzed without considering the spatiotemporally updated chemical equilibrium. Here we develop a new FRAP model aimed at long-term measurements to quantitatively determine the two distinct effects of the advection and chemical reaction, i.e., the different major sources of the change in fluorescence intensity. To validate this approach, we carried out FRAP experiments on actin in stress fibers over a time period of more than 900 s, and the advection rate was shown to be comparable in magnitude to the chemical dissociation rate. We further found that the actin–myosin interaction and actin polymerization differently affect the advection and chemical dissociation. Our results thus suggest that the distinction between the two effects is indispensable to extract the intrinsic chemical properties of the actin cytoskeleton from the observations of complicated turnover in cells.

## 1 Introduction

Subcellular multiprotein structures that shape cells are continuously replaced with surrounding proteins in cells, a phenomenon called turnover (Smith et al. 2013). Short-term turnover, which is often analyzed in cell biology studies, is driven mainly by the Brownian motion-based diffusion and chemical replacement. The diffusion-dominant turnover is typically completed within a few seconds because of the fast diffusive nature (Elowitz et al. 1999; Mullineaux et al. 2006). Meanwhile, the chemical replacement occurs over a wider range of time scales that can take several minutes (Murthy and Wadsworth 2005; Campbell and Knight 2007). The latter case, involving chemical reactions, is potentially subjected to an additional physical factor, advection, namely the transport of the proteins by local bulk motion. The involvement of advection becomes evident particularly when proteins of interest closely bind to actin cytoskeletal structures as they can be translocated due to the myosin-driven and/or actin polymerization-induced retrograde flow (Gardel et al. 2008; Yamashiro and Watanabe 2014). Thus, the interpretation of the long-term turnover can be more complicated than the short-term one because of the involvement of intracellular advection.

Fluorescence recovery after photobleaching (FRAP) is a technique widely used to evaluate the protein turnover (BLONK et al. 1993; Lorén et al. 2015). In analyzing the recovery of fluorescent proteins, a single exponential fitting is often used to reflect the kinetic rate (Campbell and Knight 2007; Dukic et al. 2017). This simple approach is often taken in the case of chemical reaction-dominant turnover as the inverse of the time constant in the single exponential function corresponds to the dissociation rate (Sprague et al. 2004). In the diffusion-dominant turnover, on the other hand, the spatiotemporal profile of the fluorescence recovery is analyzed using a diffusion equation (Crank 1975); (Braeckmans et al. 2003). In more complex cases where the diffusion and chemical reaction potentially contribute nearly equally to the turnover, elaborate physicochemical models have been used to separately characterize the two distinct factors (Sprague et al. 2006; Mueller et al. 2008). We also previously developed a platform based on FRAP analysis that allows, for the first time to our knowledge, to determine the protein domain-level chemical properties and pure diffusion coefficient of actin binding proteins (Saito et al. 2021).

Importantly, to our knowledge, all previous FRAP models have assumed that the chemical equilibrium distribution is spatially identical and immobile during the measurement. The analysis has consequently been limited to short-term observations where intracellular advection and resulting spatial shift of the actually “ mobile” scaffold proteins are ignored so that the above assumptions remain valid. To assess, instead, long-term FRAP behavior where the single exponential fitting is irrelevant, a double exponential model was alternatively used (Campbell and Knight 2007; Sakurai-Yageta et al. 2015). However, the relevance of the two independent time constants determined in the model to the actual physical factors, diffusion, chemical reaction, and/or spatial advection, is unclear.

Here we present a new model to evaluate the long-term turnover involving both the chemical reaction and advection in FRAP analyses. To demonstrate this, we analyze β-actin associated with stress fibers (SFs), a cellular contractile apparatus composed mainly of nonmuscle myosin II (NMII) as well as actin. According to previous studies with single/double exponential fitting approaches (Campbell and Knight 2007), the turnover of actin in SFs takes a time constant of ∼5 min, and the bleached region recovers spatially uniformly. The actin turnover is thus suggested to be dominated by chemical replacement rather than diffusion (Saito et al. 2021). The relatively long time constant of minutes also suggests that proteins in bleached region are susceptible to advection-driven transport. To decouple the two coexisting factors, chemical reaction and mechanical advection, in interpreting the FRAP data, here we describe an advection-reaction equation to determine, with the spatiotemporal recovery profiles, the chemical dissociation rate between actin and the scaffold SFs as well as the advection velocity. Our approach, thus appropriately modifying the bleach intensity distributions, provides a platform allowing for FRAP analyses on spatial advection-associated long-term turnover.

## 2 Materials and methods

### 2.1 Model description

We consider the β-actin turnover in single SFs where chemical reaction is dominant. In a long-term case, the fluorescence intensity in FRAP experiments will not only recover due to the chemical replacement but also change because of advection-induced spatial shift of the bleached region. A specific example shows that the intensity, which is averaged over the bleached region as typically done in single/double exponential fitting approaches, increases predominantly by advection in the middle of the observation (Fig. 1). To address these situations accompanied by advection, the time rate of change of concentration of actin bound to a single SF, *C*, is described by

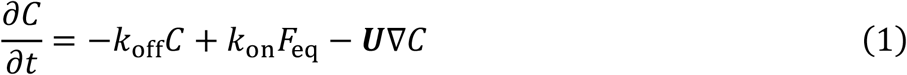

where *F*_eq_ represents the concentration of free actin molecules at unbound state, *k*_off_ and *k*_on_ represent the dissociation and association rate, respectively, *U* represents the velocity vector of the advection, ∇*C* represents the gradient, and *t* is time. To obtain an analytical solution from this advection–reaction equation, we first consider the following chemical reaction-dominant equation

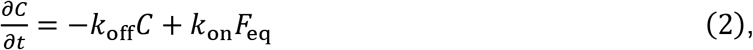

and its solution is described by

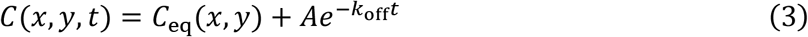

where *C*_eq_(*x, y*) represents the concentration profile at the pre-bleach state assumed to be the equilibrium state, *x* and *y* are the spatial coordinate, and *A* is the arbitrary constant determined by the initial condition. Describing the initial condition as *ϕ*(*x, y*), the solution of eq. (2) is now shown as

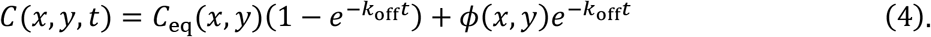

**Figure 1.**
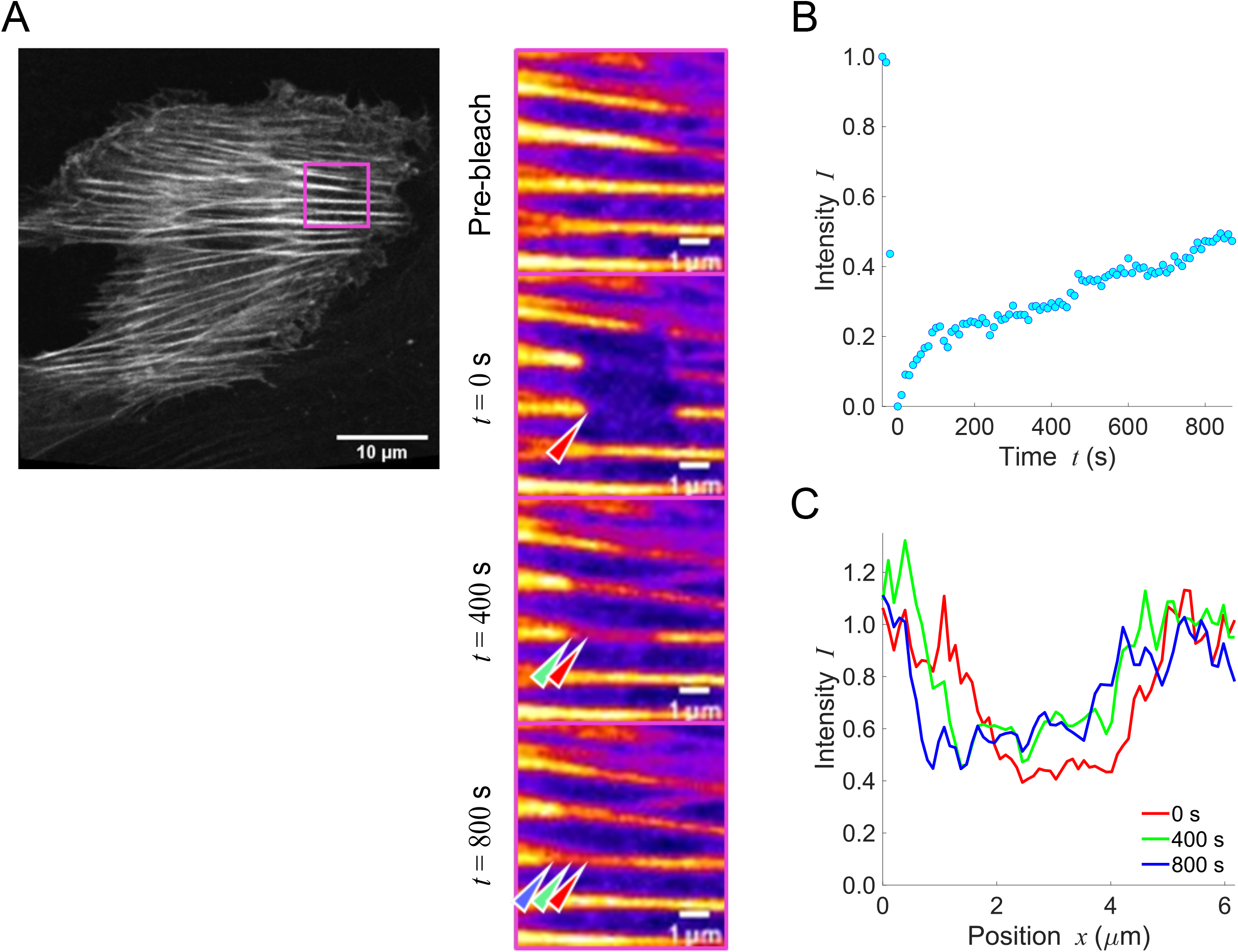
The presence of intracellular advection can impair the measurement of protein properties in long-term FRAP experiments. **A** The rectangular region in the left image (mClover2-tagged β-actin in an A7r5 cell) is magnified on the right panels, in which FRAP responses are shown. The arrow heads represent the bleached region that is being spatially shifted over time; red, green, and blue represent an identical part of the bleached rectangular region at *t* = 0, 400, and 800 s after the photobleaching, respectively. **B** Time-series change in fluorescence intensity obtained based on a conventional approach using a spatially fixed region for analysis. The FRAP response is obviously not properly approximated by the typical approach using exponential curves. **C** The intensity profiles along the length of a single SF at *t* = 0, 400, and 800 s. Scale, 10 μm.

The analytical solution of eq. (1) is now described by

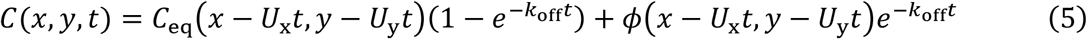

where *U*_x_ and *U*_y_ represent the velocity in *x* and *y* direction, respectively.

To determine the specific form of *ϕ*(*x, y*), first, spatiotemporal fluorescence recovery in rectangular photobleached region is described by

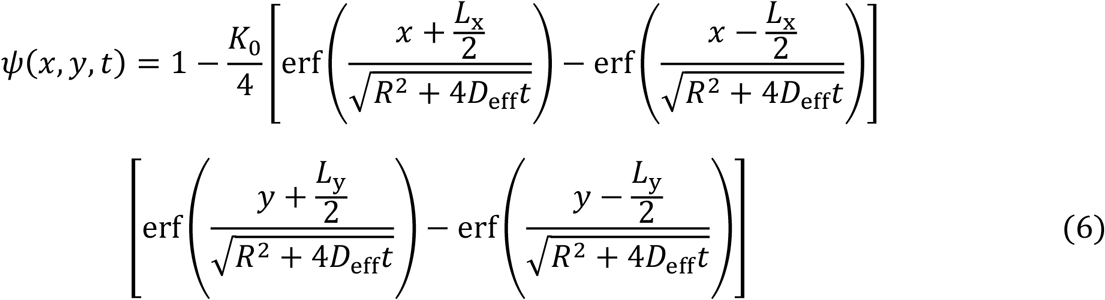

where *K*_0_, *R, L*_x_ and *L*_y_, and erf represent the photobleaching efficiency, laser resolution, width and height of the bleached rectangular area, and the error function (i.e., 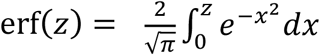), respectively (Deschout et al. 2010); and, *D*_eff_ represents the effective diffusion coefficient defined as *D*_eff_ = *D*_pure_/(1 + *K*^−1^) where *D*_pure_ and *K* represent the pure-diffusion coefficient of free actin molecules at unbound state and equilibrium dissociation constant, respectively (Crank 1975; Sprague et al. 2004; Ait-Haddou et al. 2010; Saito et al. 2021). As we previously demonstrated (Saito et al. 2021), actin in SFs is subjected to slow turnover and is predominantly at bound state, resulting in that *K* ≪ 1, and consequently *D*_eff_ ≈ 0. This drop of *D*_eff_ is reasonable because actin intensity in bleached region indeed recovers spatially uniformly in FRAP experiments (Supplementary Fig. S1). Now, eq. (6) is reduced to

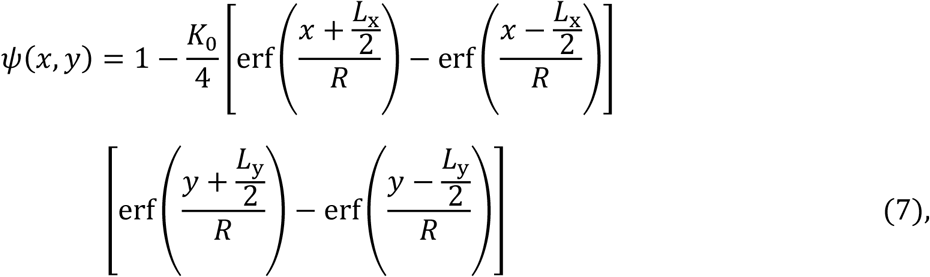

indicating that *ψ*(*x, y*) = 0 for −*L*_x_/2 < *x* < *L*_x_/2 and −*L*_y_/2 < *y* < *L*_y_/2 that correspond to bleached region, while *ψ*(*x, y*) = 1 at the outside. Next, focusing on a rectangular region of interest that contains a single SF running parallel to the long axis (*x*) of the rectangle, we assume that *C*_eq_(*x, y*) is described with error functions based on eq. (7) by

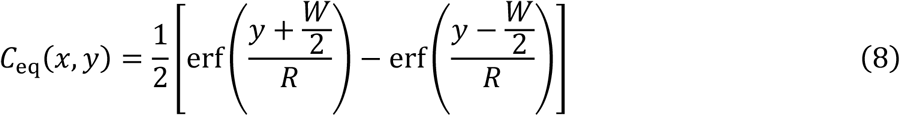

where *W* represents the width of the individual SF (Fig. 2A); here, *C*_eq_(*x, y*) = 1 for arbitrary *x* and −*W*/2 < *y* < *W*/2 where the SF is present, while *C*_eq_(*x, y*) = 0 at the other region. Finally, the initial condition is described as the product of the equilibrium distribution and photobleaching distribution (Fig. 2B):

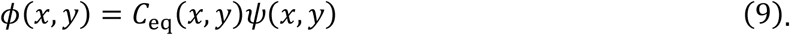

**Figure 2.**
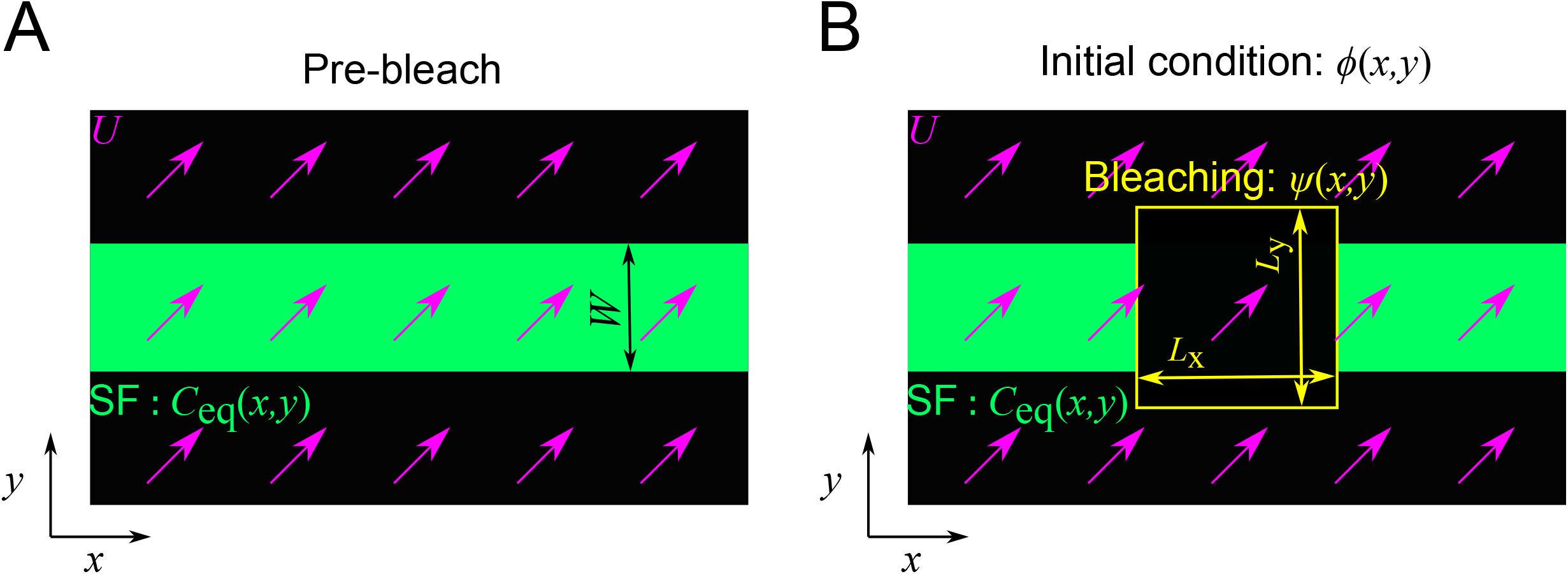
Schematic of the equilibrium distribution and initial condition in the numerical model. **A** The equilibrium distribution (green) corresponds to the intensity distribution of a single SF. Purple arrows represent the velocity vectors. **B** The initial condition is expressed by the combination of post-bleach intensity distribution (yellow) and the equilibrium distribution (green).

Accordingly, the time evolution of the advection–reaction turnover is now described by eq. (5) with the measured pre-bleach intensity distribution.

### 2.2 Numerical analysis for model validation

To demonstrate the validity of the model, we performed continuum simulation of the advection–reaction turnover. We simply consider a square region subjected to uniform advection with the same velocity *U* in both x and y directions; consequently, ∂/∂x = ∂/∂y, and *U*_x_ = *U*_y_ = *U*. By defining *t*^*^ = *k*_off_*t* and *x*^*^ = *x*/*L*_sf_ where *L*_sf_ is the longitudinal length of a single SF as a dimensionless time and length, respectively, eq. (1) of the advection–reaction equation is rewritten by

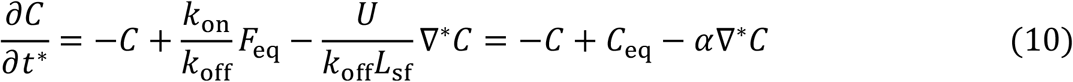

where ∇^*^*C* represents the dimensionless gradient of intensity, and *α* represents the ratio of the time for spatial shift to that for the chemical dissociation, namely

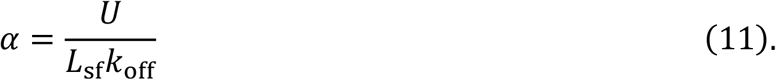

Note that *α* = 0 indicates completely chemical reaction-driven turnover. For numerical analysis, eq. (10) is discretized by the first-order upwind scheme (Supplementary materials) and calculated with eq. (9) as the initial condition. The intensity distribution is fitted by the least-square method with eq. (5). Parameters used were summarized in Table 1.

**Table 1.** Parameters used for the numerical analysis in Figure 3.

|  | Parameter | Value |
| --- | --- | --- |
| Width of SF | $W$ | $0.20L_{\text{sf}}$ |
| Bleaching length | $L_x$ | $0.50L_{\text{sf}}$ |
| Bleaching width | $L_y$ | $0.50L_{\text{sf}}$ |
| Resolution | $R$ | $0.021L_{\text{sf}}$ |
| Bleaching efficiency | $K_0$ | 1.0 |

### 2.3 Cell culture, plasmids, transfection, and inhibitors

Rat aortic smooth muscle cell lines (A75r, ATCC) were cultured with low-glucose (1.0 g/L) Dulbecco’s Modified Eagle Medium (Wako) containing 10% (v/v) heat-inactivated fetal bovine serum (SAFC Biosciences) and 1% penicillin-streptomycin (Wako) in a humidified 5% CO2 incubator at 37□. Expression plasmids encoding mClover2-tagged β-actin were constructed by inserting a human β-actin gene, which was digested with XhoI and BamHI restriction enzymes from the EYFP-actin vector (#6902-1, Clontech) into the mClover2-C1 vector (Addgene plasmid #54577, a gift from Michael Davidson). The plasmids were transfected to cells using Lipofectamin LTX with Plus Reagent (Thermo Fischer Science) according to the manufacturer’s instruction. Myosin II ATPase inhibitor (-)-blebbistatin (Wako) was used at 10 μM concentration, and actin polymerization inhibitor Latrunculin A (Wako) was used at 10 nM concentration. An equivalent amount of dimethyl sulfoxide (DMSO) was administered to cells as vehicle control.

### 2.4 FRAP experiments

Cells were cultured on a glass-bottom dish and transfected with the plasmid for 24 h. FRAP experiments were performed by using a confocal laser scanning microscope (FV1000, Olympus) with a 60x oil immersion objective lens (NA = 1.42). FRAP images were acquired every 10–15 s for 15–30 min using a 488-nm wavelength laser. The pre-bleach images were acquired for 1–2 frames before the photobleaching. A square region of 40 × 40 pixels (corresponding to *l*_x_ and *l*_y_, respectively) containing a single SFs was bleached by using 405-nm and 440-nm wavelength lasers for 1–2 frames after the pre-bleach images were acquired. For conventional FRAP analysis, images were analyzed by using ImageJ (NIH) and MATLAB (MathWorks). The time evolution of the fluorescence intensity was spatially averaged over the bleached area and was normalized by the average intensity of the pre-bleach frames.

### 2.5 Advection-reaction model-based FRAP analysis

To extract individual SFs and determine their width *W*, we set a certain fluorescence threshold using ImageJ. MATLAB was used for the subsequent data analysis. Specifically, the fluorescence intensity distribution was normalized by the spatial and temporal average of the target SF in the pre-bleach frames. The spatiotemporal evolution of the normalized intensity distribution was then fitted by the least-square method with eq. (5) to determine the parameters *θ* = (*K*_0_, *k*_off_, *U*_x_, *U*_y_).

### 2.6 Orientation analysis

The relationship between the orientation of individual SFs and the direction of advection velocity was quantified by using ImageJ. To this end, the orientation index, defined as the angle between the orientation vector of a SF and velocity vector, was obtained by calculating

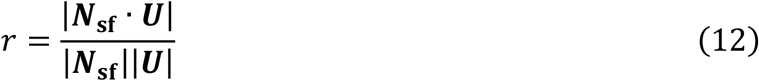

where ***N***_sf_ and ***U*** represent the orientation vector and velocity vector, respectively. Thus, *r* = 0 and *r* = 1 indicate that the velocity vector is perpendicular to and parallel with the SF, respectively.

### 2.7 Statistical analysis

Unless otherwise stated, data are expressed as the mean ± standard deviation of more than 3 independent cells. Differences were calculated based on the Mann-Whitney U-test for variables with a non-Gaussian distribution, with a significance level of *p* < 0.05 (*) or *p* < 0.01 (**).

## 3. Results

### 3.1 Numerical simulation validates the advection-reaction model

We numerically analyzed the advection-reaction turnover in FRAP using eq. (10) to evaluate the analytical solution of eq. (5). The characteristic parameter *α* was set to 0.1, 0.5, 1.0, or 10.0. Note again that *α* = 0 indicates a chemical reaction-dominant case with no spatial shift. The intensity distribution at the pre-bleach state (i.e., at the equilibrium state) was defined by eq. (8) (Fig. 3A). The time course of the intensity distribution was computationally obtained for each *α* case with the initial condition defined by eq. (9) (Fig. 3B; Video S1). The analytical solution of eq. (5) was then used to fit the numerical result by the least-square method (Fig. 3C; Video S2) to estimate the equivalent of *α*. The difference between the simulation and analytical solution in *α* was only within 2% (Table 2), suggesting that the analytical advection-reaction model well captures the turnover process undergoing spatial shift.

**Table 2.**
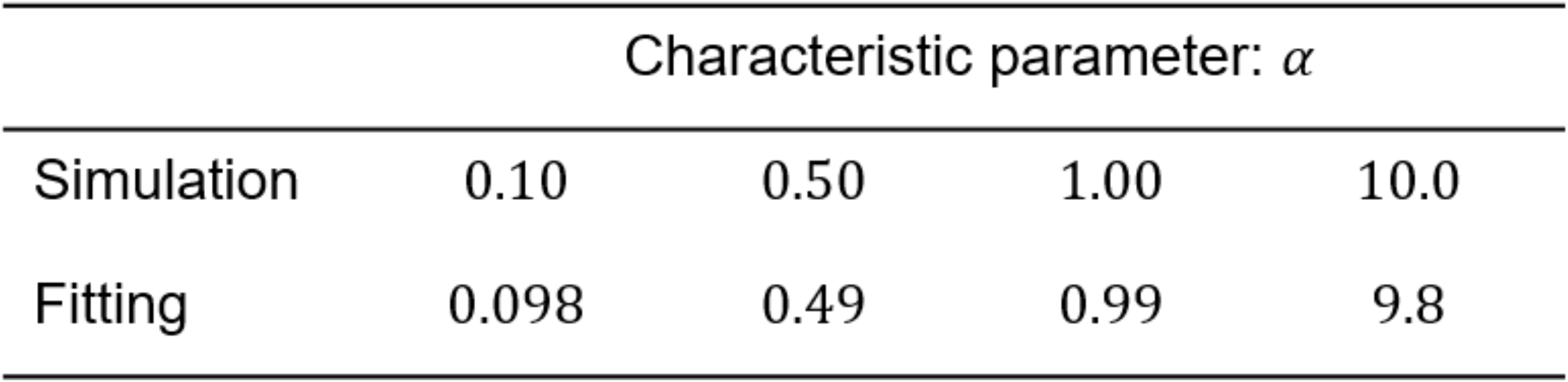
Characteristic parameter *α* used for the numerical analysis and determined by the least-square method in Figure 3.

**Figure 3.**
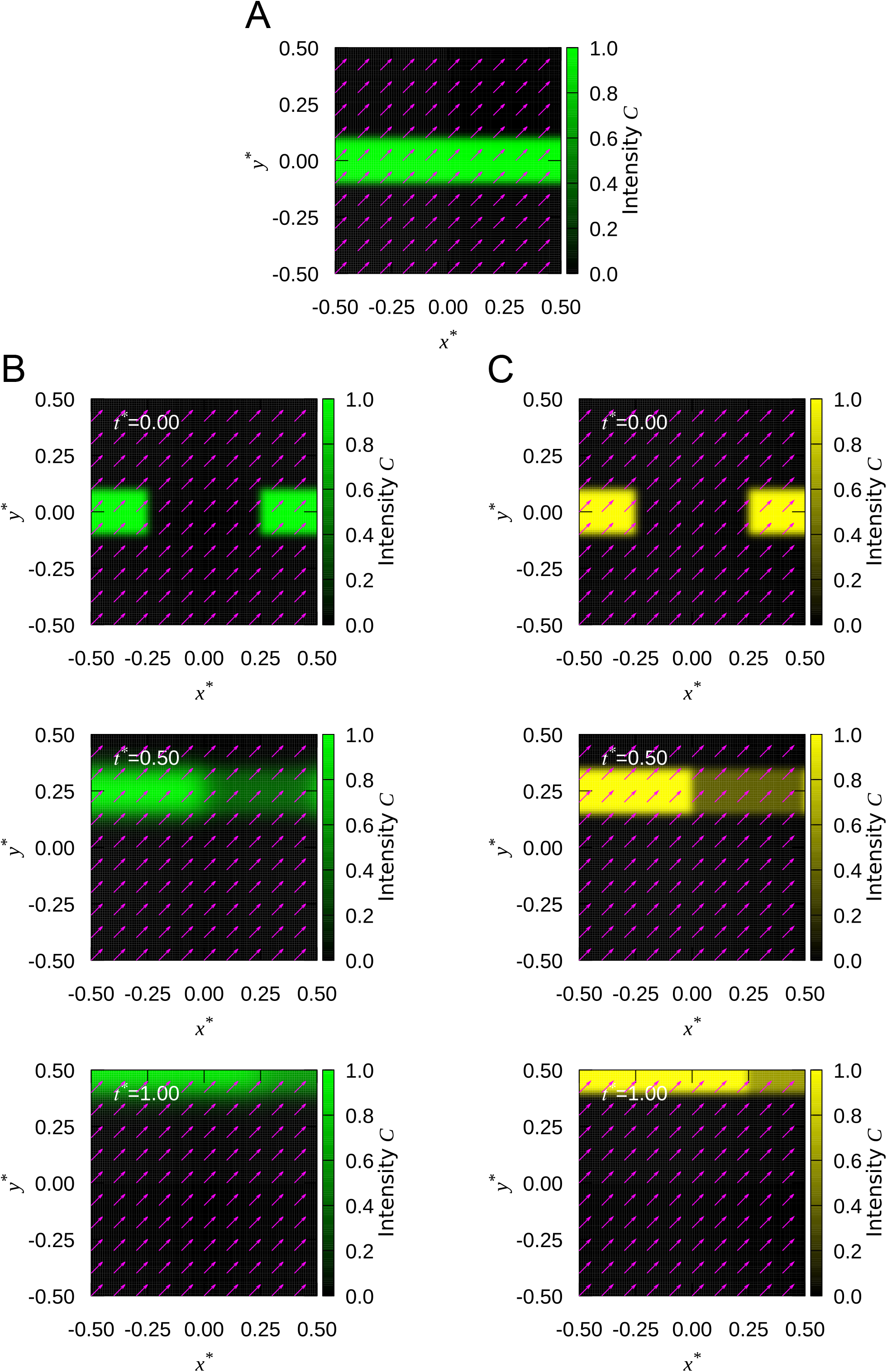
Numerical simulation validates the advection-reaction model. **A** The pre-bleach distribution expressed by eq. (8). The arrows represent the velocity vectors. **B** FRAP responses simulated by the first-order upwind scheme (see Video S1). **C** FRAP responses fitted by the least-square method (see Video S2).

### 3.2 Advection-reaction model quantitatively decouples the two distinct effects

Multiple rectangular regions were simultaneously photo-bleached in identical cells (Fig. 4A). Focusing on individual SFs, the time evolution of the intensity distribution was obtained (Fig. 4B; Video S3). The turnover of actin molecules allowed the intensity profile within the bleached region to increase over time. Meanwhile, the intensity distribution was obviously spatially shifted due to the local advection. The intensity distribution was then spatiotemporally fitted with eq. (5) (Fig. 4C; Video S4), decoupling the two distinct contributions to the fluorescence recovery, namely the chemical reaction-driven turnover and the advection. The intracellular advection is visualized by arrows representing the local velocity. The dissociation rate that characterizes the chemical reaction was determined to be *k*_off_ = (2.6 ± 2.1) × 10^−4^ s^-1^. The magnitude of the velocity was |*U*| = (5.9 ± 4.1) × 10^−4^ μm/s. Assuming that the characteristic length is on the order of 1 − 10 μm given the order of the axial length of the individual SFs within cropped images, it turns out that *α* = *U*/*L*_sf_*k*_off_ ∼0.2 − 2. This estimate suggests that the velocity of the spatial shift was comparable in magnitude with or only one order of magnitude smaller than the dissociation rate, justifying that the effect of the spatial shift or advection is not negligible.

**Figure 4.**
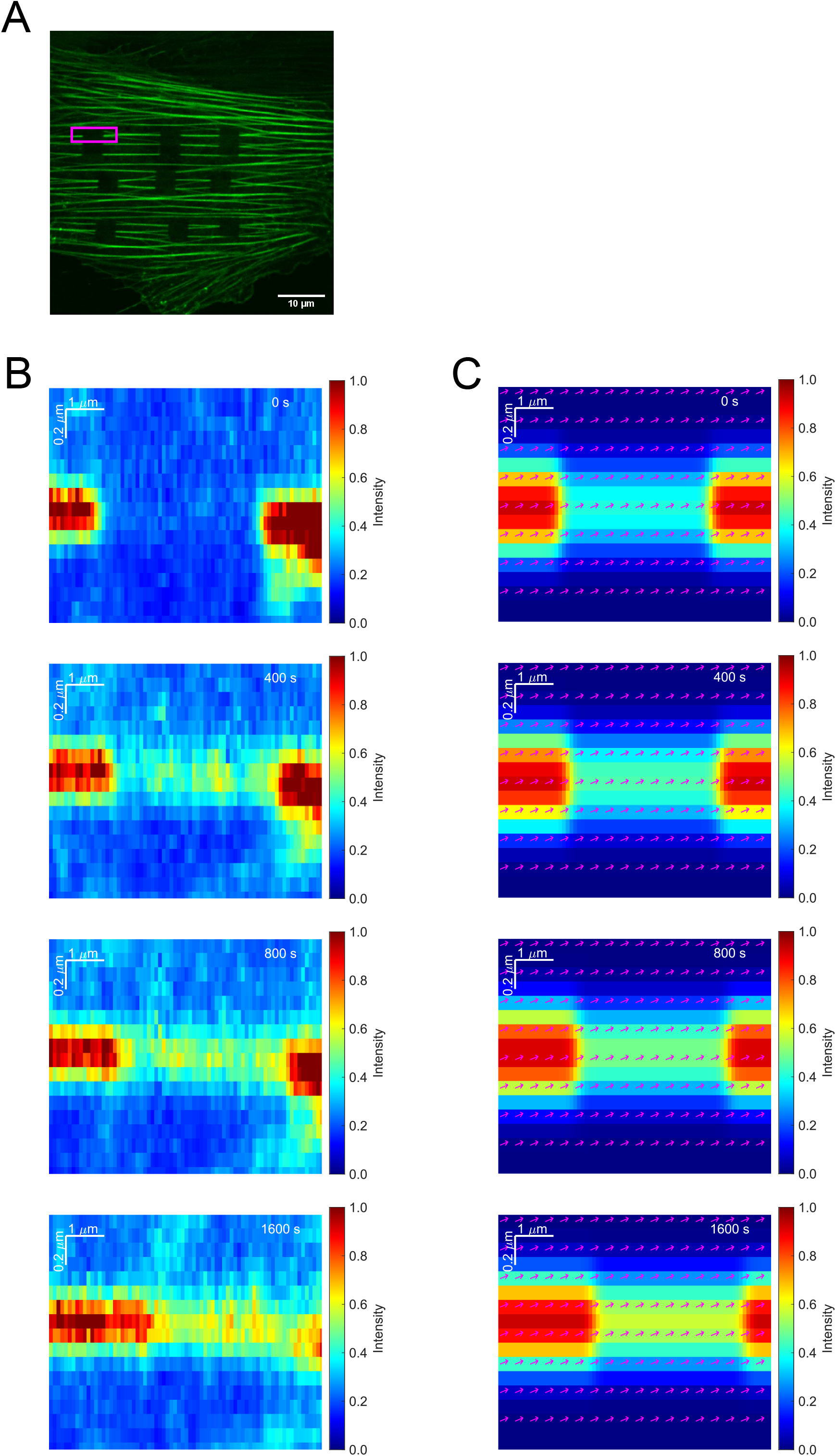
The advection-reaction model fitted to experimental data. **A** FRAP was performed onto SFs in an A7r5 cell, in which mClover2-tagged β-actin is expressed. **B** Experimentally obtained two-dimensional FRAP response of a single SF (rectangular region in A; see Video S3). **C** The model was fitted with the least-square method to the spatiotemporal evolution of the experimentally obtained FRAP response (see Video S4). The arrows represent the velocity vectors. Scale, 10 μm.

### 3.3 Actin polymerization and myosin II activity contribute differently to the advection and reaction

FRAP experiments were performed on cells treated with a moderate concentration of (-)-blebbistatin (Blebb, 10 μM), latrunculin A (LatA, 10 nM), or its vehicle DMSO (Fig. 5A-5C). Here, Blebb is an inhibitor of the phosphate release in the NMII’s ATPase cycle, and LatA is an inhibitor of actin polymerization. The parameters *θ* = (*K*_0_, *k*_off_, *U*_x_, *U*_y_) were determined by fitting eq. (5), with the least-square method, to the time-series data of the spatiotemporal distribution of the intensity. We then obtained the dissociation rate of actin from SFs (Fig. 5D) and the advection velocity (Fig. 5E). Compared to control, the dissociation rate was significantly increased with Blebb, likely because of the loss of the function of NMII as a major cross-linker to bundle the actin filaments into SFs (Matsui et al. 2011). In contrast, the dissociation rate was significantly decreased with LatA, suggesting that the decreased actin polymerization slows the dissociation of actin from SFs. The advection velocity was significantly increased in both cases of Blebb and LatA compared to control, in which LatA led to a larger increase. The orientation index quantified with eq. (12) suggests that the intracellular advection occurs predominantly in the axial direction of SFs regardless of the conditions (Fig. 5F–5H).

**Figure 5.**
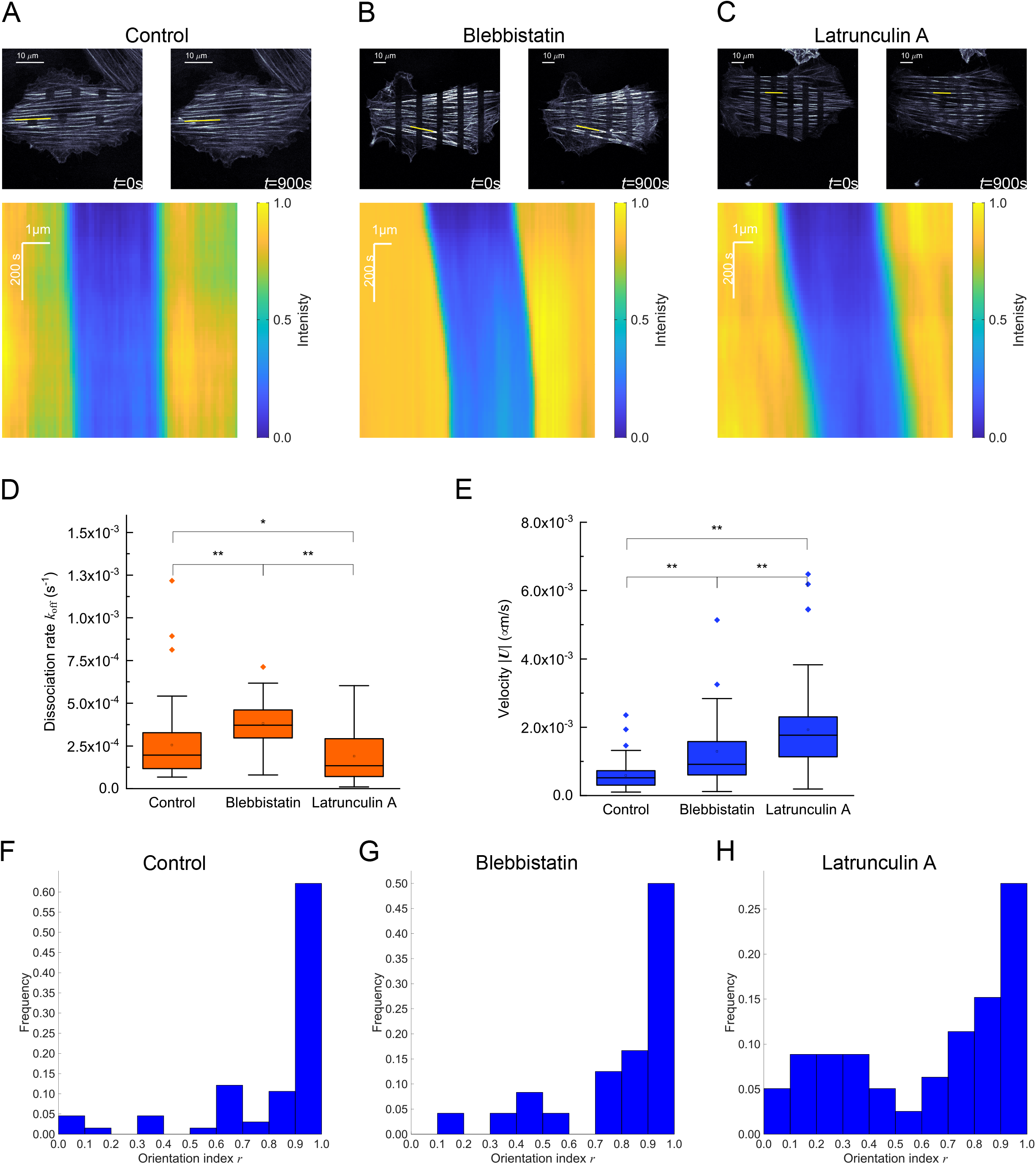
The effect of actin–myosin interaction and actin polymerization on the inherent properties of actin on SFs and intracellular advection. **A**–**C** Cells expressing mClover2-tagged β-actin and undergoing FRAP at vehicle control (**A**), Blebb (**B**), and LatA (**C**) conditions. Kymographs of the fluorescence intensity along the specified yellow lines are shown below for each condition. **D, E** The dissociation rate (**D**) and advection velocity (**E**) of actin on SFs for each condition shown by mean ± SD (*n* = 66, 24, and 52 regions of interest for vehicle control, Blebb, and LatA conditions, respectively, from at least 3 independent experiments). Box and whisker plots show the following: the upper and lower edges of the boxes represent the 75 and 25 percentile ranges, respectively; the central lines represent the median; the whiskers represent the standard deviation; the open dots represent the mean; and, the closed dots represent outliers. The asterisks indicate significant differences. **F**–**H** The histogram of the orientation index that describes the correlation in the directions of SFs and advection velocity vectors for vehicle control (**F**), Blebb (**G**), and LatA (**H**) conditions (*n* = 66, 24, and 52 regions of interest for vehicle control, Blebb, and LatA conditions, respectively, from at least 3 independent experiments). Scale, 10 μm.

## 4. Discussion

A number of physicochemical models have been developed to analyze the FRAP behavior in diffusion-dominant, chemical reaction-dominant, or their mixed cases (Lorén et al. 2015). In a chemical reaction-dominant case, the turnover typically occurs on a time scale of minutes. Particularly for longer-term observations, another factor namely advection inevitably affects the apparent recovery of the fluorescence intensity to consequently impair the accuracy of the measurement (Fig. 1). To our knowledge, however, no FRAP model has allowed for decoupling the mixed effects contaminated by such bulk flow-like movements or spatial shifts of target molecular complexes. We then developed a new FRAP model based on the advection-reaction equation and demonstrated the simultaneous determination of the separate effects by applying it to the actin cytoskeleton on SFs. We determined the chemical dissociation rate and the advection velocity from data taken for a long period of time over 900 − 1800 s, which must have not been achieved previously. Our results showed that the characteristic parameter *α* = *U*/*L*_sf_*k*_off_ ∼0.2 − 2, suggesting that the intracellular advection occurs on a time scale comparable to the dissociation of actin from SFs, and thus that the distinction between the two effects is indispensable to accurately evaluate their inherent properties within cells.

The intracellular advection or retrograde flow is created due to the actin–myosin interaction as well as actin polymerization, both of which also drive the generation of SFs (Hotulainen and Lappalainen 2006). The consistency of the main direction of the advection with that of SFs, we detected here (Fig. 5F–5H), seems therefore reasonable. To probe the individual effects of the two distinct factors on the extent of actin dissociation and advection, we treated cells with Blebb and LatA. With Blebb treatment that inhibits the actin–myosin interaction, the dissociation of actin from SFs was enhanced (Fig. 5D). This tendency is consistent with the view that NMII works as a major cross-linker to bundle actin filaments into SFs, in which the loss of the actin–myosin interaction causes the unbundling of SFs into individual actin filaments (Matsui et al. 2011; Okamoto et al. 2020; Saito et al. 2020; Huang et al. 2021) (Fig. 6). Thus, the increased turnover in the FRAP response is likely to be caused at the level of actin filaments but not that of actin monomers.

**Figure 6.**
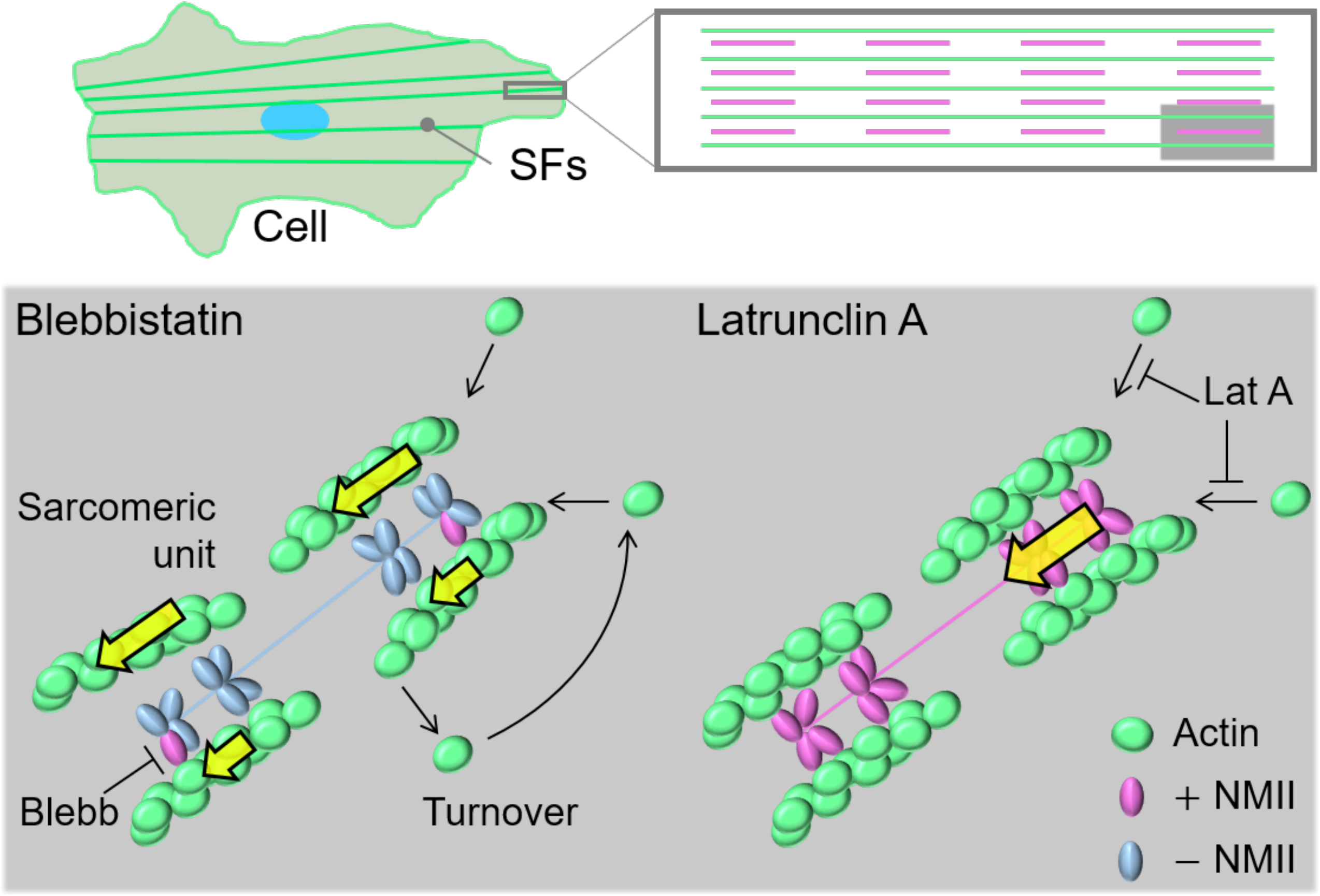
Schematic to illustrate the actin turnover in the sarcomeric unit of SFs at the different drug treatment conditions. With Blebb treatment, the entire actin filaments are dissociated from the sarcomeric unit to increase the actin dissociation rate. Meanwhile, the unbundled and thus movable actin filaments may now contribute to the increase in the advection velocity (yellow arrows). With LatA treatment, the resulting increase in actin monomers in the cytoplasm may slow the depolymerization of preexisting actin filaments and thus decrease the actin dissociation rate. Meanwhile, the sarcomeric unit tends to slide quickly (yellow arrows) potentially because of the maintained NMII activity, partly damaged physical interface at the adhesions, and activated lamellipodia/filopodia-driven actin motility.

On the other hand, the dissociation of actin in SFs is reduced upon the inhibition of actin polymerization (Fig. 5D), suggesting that apparently it rather stabilizes the preexisting actin filaments. This result seems to be in contradiction with the observations that 30-min LatA treatment finally increases the ratio of actin monomers to actin filaments within cells (Huang et al. 2021). At the level of the sarcomeric unit or contractile unit of SFs, however, the absence of new addition of actin monomers may slow the depolymerization of the constituent actin filaments into actin monomers for some time, relaxing their release to the increasing pool of actin monomers in the cytoplasm.

Together with these inherent responses of actin in SFs, the velocity field of intracellular advection was simultaneously obtained in our analysis. The mean velocity was significantly increased upon the treatment with Blebb (Fig. 5E), which is consistent with previous reports (Liu et al. 2010; Nehwa et al. 2020). As there is a competitive signaling relationship between Rho-mediated formation of the mature actin structures of SFs and Rac1/Cdc42-mediated formation of more dynamic actin structures of lamellipodia and filopodia (Nobes and Hall 1995), the limited activity of SFs stimulates the lamellipodia/filopodia-associated actin motility, which may in turn increase the motility throughout the whole cells where the unbundled and thus movable actin filaments are increasing in number (Fig. 6). Besides, the intracellular flow is known to be enhanced with the deterioration of the physical connection of SFs to the cell– substrate adhesions (Oakes and Gardel 2014), which may also be caused by Blebb treatment given the decreased cellular forces to be exerted at the substrate (Nehwa et al. 2020).

The advection velocity was more significantly increased upon the treatment with LatA (Fig. 5E). To gain insights on this behavior, we examined the effect of treating cells with a higher LatA concentration of 10 μM. We then observed a severing of SFs at their termini where cell–substrate adhesions are located and their subsequent extensive retraction (Video S5), suggesting that the complete loss of the supply of new actin monomers is fatal to the anchoring of SFs to the substrate, whereas their contractility is still kept active. This observation led us to propose that, because of the less structural integrity due to the limited actin incorporation compared to control, the interface of SFs to the adhesions tend to disintegrate, and hence the other more intact parts of the identical SFs slide faster as they are subjected to the NMII-driven retrograde flow while substantially keeping the sarcomeric structure (Fig. 6). The advection velocity will consequently be increased because of the faster sliding compared to control, while the turnover of actin is rather suppressed as already discussed above.

In summary, here we described a new FRAP model, which considers the effect of intracellular flow-induced spatial shift of target materials and thus is critical to long-term measurements where the shift is no longer negligible. We demonstrated that, while conventional approaches with a spatiotemporally fixed region of analysis fail to accurately determine the inherent chemical constant of actin on SFs, the present analysis allows for determining it as well as the spatiotemporal evolution of the advection field. Using this model, we analyzed how the inhibition of actin–myosin interaction and actin polymerization separately affect the actin dynamics on SFs. The response to these molecular perturbations was discussed at the level of the sarcomeric unit. As the integrity of SFs has been implicated in many signaling events including the activation of proinflammatory pathways (Deguchi et al. 2006; Kaunas et al. 2010), the methodologies and findings obtained here will be useful particularly in evaluating the inherent properties of the actin cytoskeleton associated with SFs.

## Acknowledgments

TS is supported by Japan Society for the Promotion of Science (JSPS). This study was partly supported by JSPS KAKENHI Grants (18H03518, 19K22967, and 20J10828).

## Data availability

The data that support the findings of this study are available from the corresponding author upon reasonable request.

## Supplementary materials

### The first-order upwind scheme for the numerical calculation

To numerically calculate the advection-reaction equation, the first-order upwind scheme was used as follow:

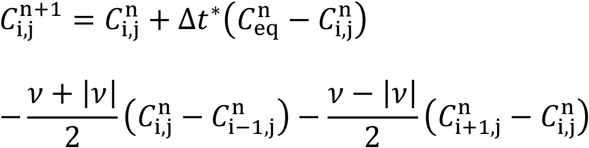

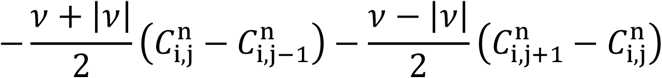

where 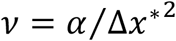, and n, i, and j represent the iteration of calculation, position along x, and that of y, respectively.

**Figure S1.**
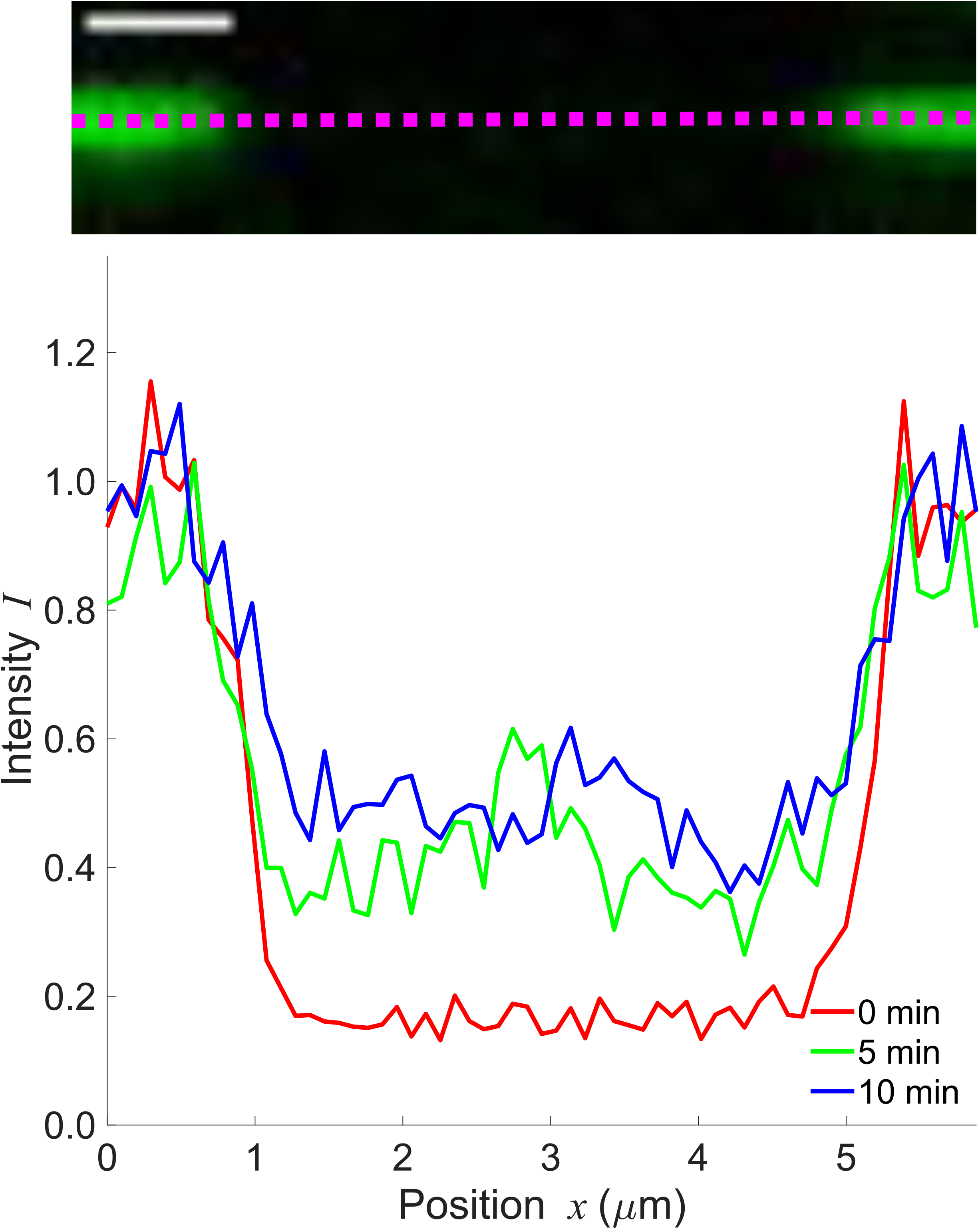
Time-course of line profile along a SF, indicating spatially uniform recovery in the bleached region (without diffusive gradient). Scale, 1 μm.

Video S1 FRAP response demonstrated by the numerical analysis.

Video S2 Numerically analyzed FRAP response (Video S1) was fitted with the model by the least-square method.

Video S3 FRAP images in a single SF.

Video S4 FRAP images (Video S3) were fitted with the model by the least-square method.

Video S5 mClover2-tagged β-actin in a cell subjected to 10-μM LatA.

